# Physiological Processing of Aversiveness in the Mind’s Ear: Comparing Audiovisual, Auditory and Imagined Auditory Affective Stimuli

**DOI:** 10.1101/2024.04.29.591662

**Authors:** Xuan Yang, Sewon Oh, Jacob Stanley, Sarah Hammons, Ashley Anderson, Douglas H. Wedell, Svetlana V. Shinkareva

## Abstract

Understanding the somatovisceral responses to auditory affective imagery has important implications in disorders like misophonia. The current study compared physiological responses to aversive and nonaversive states across three modalities: audiovisual, auditory, and auditory imagery. Electromyographic activity over corrugator supercilii (EMGc) and zygomaticus major (EMGz), electrodermal activity (EDA), heart rate (HR), and finger skin temperature (SKT) were measured. There was significant differentiation in EMGc, EDA, and HR deceleration between aversive and nonaversive audiovisual stimuli. EMGc potentiation was the only physiological measure showing consistent differentiation across the three modalities. Cross-modal aversiveness classification results revealed a similar physiological response pattern between audiovisual and auditory modalities. The physiological response pattern during auditory affective imagery was useful for predicting the aversiveness in audiovisual modality but not the other way around. Vividness in auditory imagery correlated with subjective hedonic valence ratings, but not physiological responses. Taken together, the current data suggest that the aversiveness of auditory imagery is differentiable in subjective affective experience and facial muscle potentiation. These results of the physiological responses to imagined aversive sounds in nonclinical population would serve as a comparison baseline for the study of misophonia.

## Introduction

Recognizing and processing aversiveness is vital for the survival of many organisms. In humans, aversive stimuli evoke systematic and coherent changes in both subjective experience and somatovisceral responses. Maladaptive behavioral and physiological responses to aversive stimuli have been linked to psychopathological symptoms such as depression, anxiety, and phobia (Holmes & Mathews, 2010; Lang & Cuthbert, 1984; Lang & McTeague, 2009; Rottenberg et al., 2005). Examination of the somatovisceral correlates of aversive stimuli across a variety of perceptional modalities can contribute to a better understanding of aversiveness processing across different contexts and behavioral responses.

Previous studies have shown that aversive and appetitive states can be reliably differentiated through electromyographic, electrodermal, and cardiovascular measures. Electromyogram (EMG) activity over corrugator supercilii (EMGc), a facial muscle related to brow-lowering, has been considered a gold standard measure for aversiveness. EMGc potentiates more for aversive stimuli compared to nonaversive stimuli (Bradley et al., 2001; Bradley & Lang, 2000; Brown & Schwartz, 1980; Codispoti et al., 2001, 2008; Lang et al., 1993; Sato & Kochiyama, 2022; Schwartz et al., 1980). EMG activity over zygomaticus major (EMGz), a facial muscle related to lip-pulling, has a closer correspondence to pleasure (Bernat et al., 2006; Codispoti et al., 2008; Dellacherie et al., 2011; Grewe et al., 2007). However, evidence has revealed a quadratic relationship between EMGz and valence, i.e., this muscle potentiates for extreme valenced events without differentiating between positive and negative valence (Bradley & Lang, 2000; Lang et al., 1993; Larsen et al., 2003). Electrodermal activity (EDA) reflecting the activity in sweat glands innervated by the sympathetic nervous system has a closer correspondence to arousal rather than valence. The accentuated skin conductance in response to both pleasant and unpleasant arousing stimuli has been reported from various stimulus types and body sites (Bradley et al., 2001; Bradley & Lang, 2000, 2007; Codispoti et al., 2001; Palomba et al., 2000; Sato et al., 2020; Van Dooren et al., 2012). Heart rate is innervated by both sympathetic (heart rate acceleration) and parasympathetic nervous systems (heart rate deceleration). An initial heart rate deceleration following aversive stimulus presentation has been consistently observed under varying aversive contexts, including viewing pictures (Lang et al., 1993), hearing sounds (Bradley & Lang, 2000), and watching films (Baldaro et al., 2001; Bradley et al., 2001; Palomba et al., 2000). The accentuated skin conductance and heart rate deceleration during processing of aversive compared to nonaversive stimuli may suggest a heightened sensory intake and orienting at the initial stage of the defensive motivational system activation (Bradley et al., 2001; Bradley & Lang, 2007). Skin temperature (SKT) is another marker for arousal, indexing the sympathetic induced changes in microcirculation (Kistler et al., 1998). Decreases in skin temperature due to reduced skin blood flow from cutaneous vasoconstriction have been associated with both arousing aversive and appetitive states (Ioannou et al., 2014; Levenson et al., 1991; Sato et al., 2020; Sato & Kochiyama, 2022; Sinha, 1996) and are similar to EDA in differentiating arousal levels(Sato & Kochiyama, 2022).

Evidence from the past 50 years has suggested that the somatic and autonomic responses not only corresponded to exogenously induced aversiveness but also endogenously induced aversiveness from affective/emotional imagery. Based on the Bio-Informational Theory proposed by Lang (1979), the mental imagery of affective events activates an associative network in the brain, which involves the stimulus information (the sensory and contextual description of the imagined event), the semantic information (the idiosyncratic memory deriving from previous experience or knowledge) and the response information (the somatic and autonomic responses representing real-world coping actions and reactions). Previous research has shown that physiological responses during endogenously induced affective imagery are consistent with those elicited exogenously, including potentiated facial muscle tensions (Brown & Schwartz, 1980; Schwartz et al., 1976, 1980), increased skin conductance (Acosta & Vila, 1990; Carroll et al., 1982), decreased finger skin temperature (Levenson et al., 1991; Sinha, 1996), and enlarged pupil diameter (Henderson et al., 2018). Heart rate is the only marker that showed different change patterns between external affective stimuli perception and affective mental imagery. As opposed to the sustained heart rate deceleration when viewing pictures, listening to sounds, and watching movies, imagining affective scenes have been reported to index sustained heart rate acceleration (Acosta & Vila, 1990; Carroll et al., 1982; Lang et al., 1980; Levine et al., 2016; G. A. Miller et al., 1987; M. W. Miller et al., 2002; Vrana et al., 1989), and this acceleration was further enhanced when participants imagine person-relevant scenes (M. W. Miller et al., 2002) or after imagination training (Acosta & Vila, 1990), which may suggest different underlying cognitive process (Bradley & Lang, 2007). Rather than the enhanced sensory intake from external stimuli as indexed by initial heart rate deceleration, accelerated heart rate during active imagining may reflect “action engagement prompted by the imagery scene” (Bradley & Lang, 2007).

Most investigations of affective imagery have focused on visual representation, “seeing in the mind’s eye” (see Bradley et al., 2023 for a review). The physiological responses to auditory affective imagery remain less explored. According to Lang’s Bio-informational Theory (1979), affective imagery is a construct built from stimulus-specific information, suggesting a rich, multimodal composition that goes beyond mere visual representation. Auditory imagery refers to the auditory representation of sounds in the absence of external auditory stimulus, “hearing in the mind’s ear”, which can be very vivid and closely approximate real perceptual experience (Janata, 2012; Lima et al., 2015). A deeper investigation into the physiological distinctions between aversive and nonaversive affective states in auditory imagery could extend our understanding of affective imagery from visual to auditory modality. Moreover, it could provide a comparative baseline for studying the physiological responses to aversive sounds in disorders like misophonia, an affective sound-processing disorder characterized by strong aversive emotions in response to everyday sounds (Kumar et al., 2017), and post-traumatic stress disorder.

Individual differences in vividness in visual mental imagery have been shown to modulate physiological responses of aversiveness. Miller et al. (1987) observed significant greater heart rate acceleration in good versus poor imagers when imagining fear and anger events. Wicken et al. (2021) found that aphantasia participants who report being unable to generate visual imagery did not show the increase in skin conductance that the general population exhibits when imagining frightening stories. These results support Lang’s (1979) Bio-Information Theory that vivid mental imagery consisting of rich stimulus-information would correspond to heightened physiological responses (Lang & Cuthbert, 1984). The modulation effect of imagery vividness is also found for the auditory modality. Neuroimaging studies have provided indirect evidence that auditory imagery vividness positively correlates to brain activation in higher-level auditory processing regions including the superior temporal gyrus and supplementary motor area (Halpern, 2015; Herholz et al., 2012; Lima et al., 2015). However, no study to date has examined the correspondence between auditory imagery vividness and somatovisceral responses in affective states. Empirical study is needed to examine the extent to which individual differences in auditory imagery vividness impact physiological processing of aversiveness.

To better understand somatovisceral responses to imagined aversive auditory stimuli, the current study compared responses to aversive and nonaversive states across three modalities: audiovisual (AV), auditory (A), and auditory imagery (AI). Based on previous evidence, this design allowed us to test the following set of hypotheses:

H1: Coherent facial, electrodermal, cardiac, and skin temperature changes will be elicited when processing aversive audiovisual (H1a) and auditory (H1b) stimuli. It was further predicted that heightened EMGc and EMGz potentiation will correspond to aversive and nonaversive stimuli, respectively, HR deceleration will correspond to aversive stimuli, and increase in EDA and finger SKT will correspond to both aversive and nonaversive stimuli.

H2: Similar physiological responses were predicted for auditory imagery, with the exception that acceleration, rather than deceleration, will occur in HR for aversive auditory imagery compared to nonaversive auditory imagery.

H3: Given physiological responses share a similar pattern across AV, A, and AI, an aversiveness classifier trained on the physiological signals from one modality will perform significantly better than chance level for the aversiveness classification of physiological signals from another modality.

H4: Participants who score higher on vividness of auditory imagery will show more negative valence ratings, higher arousal ratings, and heightened physiological responses for aversive compared to nonaversive auditory imagery.

## Methods

### Participants and experimental procedure

The required sample size was decided upon the *A prior* power analysis using G*Power software 3.1.9.7 (Faul et al., 2007) based on the differentiation in EMGc between aversive and pleasant visual imagery using a similar experimental design (M. W. Miller et al., 2002). A sample of more than 44 participants is needed, with α = 0.05, power (1-β) = 0.90, and an estimated Cohen’s *d* = 0.50. Fifty-nine undergraduate students (*M*_*age*_ = 20.49, *SD*_*age*_ = 2.64, 45 female) were recruited through the participant pool of the psychology department at the University of South Carolina. After signing the informed consent in accordance with the Institutional Review Board at the University of South Carolina, the participants filled out two behavioral questionnaires. They were then equipped with devices for physiological monitoring (see *physiological recording* section for details). This was followed by three practice trials, corresponding to the three presentation modalities, to ensure understanding of the experimental procedure before proceeding to experimental trials (Figure 1A).

**Figure 1.**
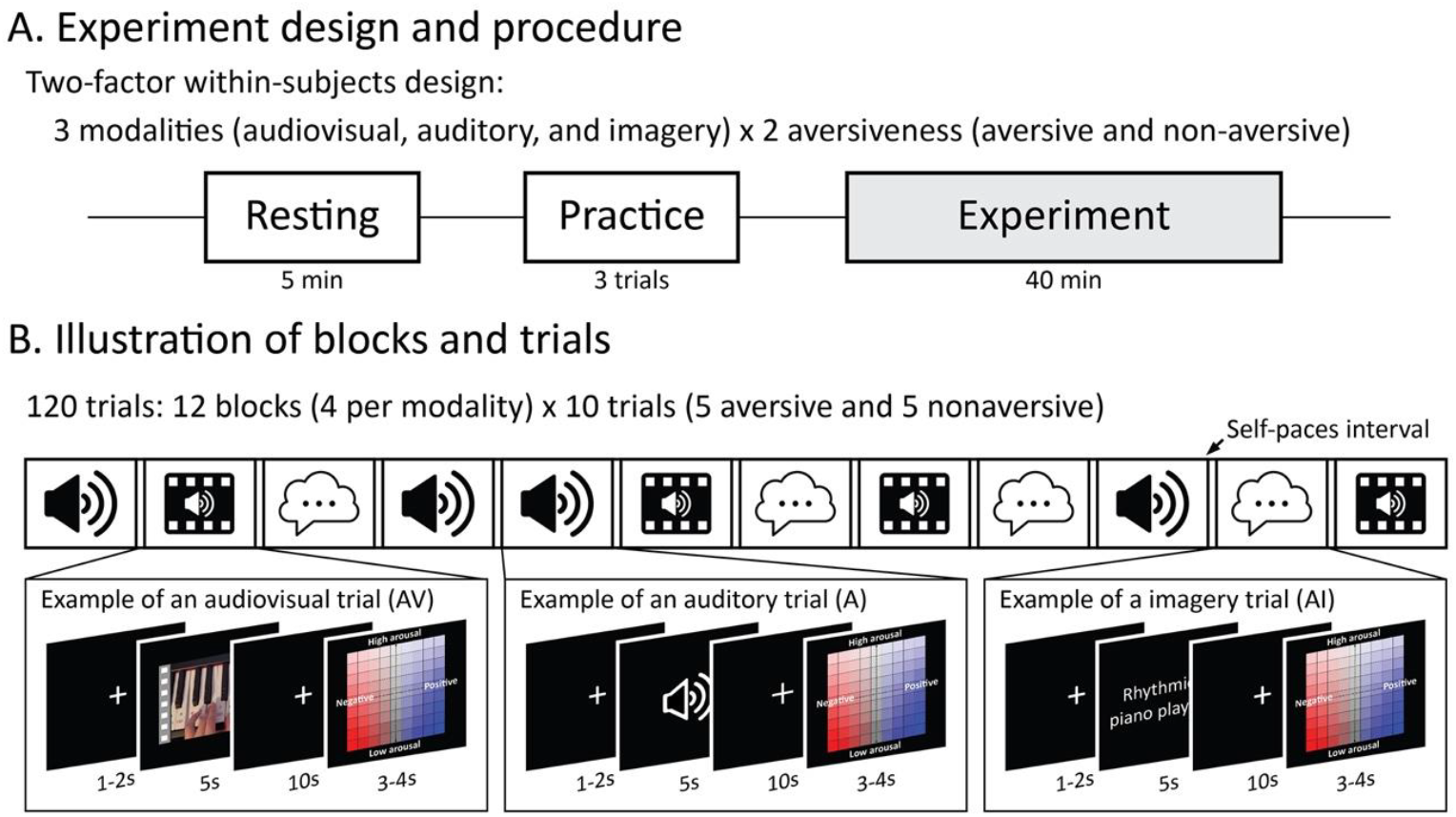
Illustration of the experiment design, including block and trial structure. A. Experiment design and procedure. B. Illustration of blocks and trials. Five aversive and five nonaversive stimuli were presented in a random order in each block, for 12 blocks. The stimuli within each block were of the same modality.

### Stimuli

The stimuli consisted of 5 s audiovisual clips that differed on aversiveness, their auditory components, and corresponding prompts for auditory imagery. We selected 20 aversive and 20 nonaversive audiovisual stimuli from an in-house database. For the auditory (A) stimuli, the audio tracks from the original audiovisual stimuli were extracted and normalized to a uniform loudness level of −16 Loudness Unit Full Scale (LUFS). These normalized audio tracks were integrated with the original videos for the audiovisual (AV) stimuli. Descriptive phrases of the stimuli that prompted participants to imagine a corresponding sound (e.g., “Rhythmic piano playing” and “Peaceful windchimes”) were used for the auditory imagery (AI) stimuli. There were 120 (20 exemplars × 2 aversiveness level × 3 modalities) unique stimuli in total.

### Experimental design

To examine the effect of aversiveness on physiological measures across modalities, we employed a 2 (Aversiveness: aversive, nonaversive) × 3 (Modality: A, AV, and AI) within-subject factorial design. For each of the three modalities, the 40 stimuli (20 aversive and 20 nonaversive) were randomly assigned to four blocks. Each block consisted of five aversive and five nonaversive stimuli, and stimuli were presented in a random order. Each trial was designed to last 20 s to ensure enough time for physiological responses’ activation and recovery. Each trial began with a 1-2 s fixation cross, followed by a 5 s stimulus presentation, another fixation cross presented for 10 s, and a rating screen for 3-4 s (Figure 1B). Participants rated each stimulus on valence and arousal using a valence-by-arousal grid, with valence ranging from 0 (negative) to 9 (positive) and arousal from 0 (low) to 9 (high).

### Physiological recording

Facial muscle and autonomic responses during the experiment were recorded using a BIOPAC MP150 recorder (Biopac, Inc., Goleta, CA, USA) with a sampling rate at 1000 Hz. EMG was recorded over corrugator supercilii and zygomaticus major on the left side of the participants’ face using a BioNomadix wireless amplifier. EMGc was captured with two electrodes placed above the eyebrow (one electrode directly above the left eyebrow on the imaginary vertical line crossing the inner corner of the eye, and the other approximately 1 cm lateral to the first). EMGz was captured with one electrode placed at the center of the imaginary line joining the left cheilion and the preauricular depression and the other placed 1 cm inferior and medial to the first electrode along that imaginary line). A ground electrode for EMG was placed in the middle of the inner eyebrows. EDA was recorded with two electrodes placed on the fingertip of the index and middle fingers of the left hand using a GSR100C amplifier. ECG was recorded in a lead II configuration with one electrode placed below the right collar bone (cathode) and the second electrode placed below the lowest left rib (anode) using an ECG100C amplifier. No additional ground cable was attached considering we have concurrent EDA data collection. SKT was recorded with a sensor placed at the thumb of the left hand using a SKT100C amplifier.

### Preprocessing and data reduction

The raw EMG signals were preprocessed with a 50 Hz band-stop filter and a 20-500 Hz band-pass filter, then rectified. The area under the curve (AUC) for the 7 s time window following stimulus presentation was calculated for each trial. A log10 transformation was applied to correct the non-normal distribution. To facilitate between-subject comparison, the data was first baseline corrected by subtracting the AUC (log-transformed) for the 1s time window prior to the stimulus presentation and then converted to z-scores across trials within participants (separately for corrugator and zygomaticus).

The phasic EDA signals were calculated online by applying a 0.05 Hz high-pass filter to the tonic EDA signals. These were then smoothed offline by averaging across every 50 samples and transformed into absolute values. The AUC for the 7 s time window following stimulus presentation was calculated for each trial. The subsequent pipeline of log-transformation, baseline correction, and standardization was the same as the EMG analysis.

We used Acqknowledge software by BIOPAC to automatically detect the R-peaks in the raw ECG waveform. Motion artifacts were manually detected and corrected. The mean heat rate (HR) was calculated for the 7 s time window following stimulus presentation based on inter-beat intervals and corrected for the mean of the 1 s pre-stimulus baseline. The data was standardized across trials within participants.

The raw SKT signals were averaged across the 7 s time window following stimulus presentation, baseline-corrected for the mean of the 1 s pre-stimulus window and standardized across trials within participants.

### Behavioral measures

Vividness of Auditory Imagery: The Vividness subscale of the Bucknell Auditory Imagery Scale (Halpern, 2015; BAIS-V) was used to measure the vividness of auditory imagery. Each of its 14 items instructs participants to construct an auditory image specific to a real-life scenario. Participants were asked to rate the vividness of those auditory images using a 7-point Likert scale, with 1 representing “no image is present at all” and 7 representing “the image is as vivid as the actual sound”. Higher scores on BAIS-V indicate higher mental imagery ability on the vividness of auditory imagery. The BAIS-V questionnaire shows high internal consistency in the original study (Cronbach’s alpha = .83) and the current study (Cronbach’s alpha = .92).

Vividness of Visual Imagery: The Vividness of Visual Imagery Questionnaire (Marks, 1973; VVIQ) was used to measure visual imagery vividness. This questionnaire consists of 16 items. Participants were asked to construct a visual image specific to a scenario and rate the vividness on a 5-point Likert scale, with 1 representing “Perfectly clear and as vivid as normal vision” and 5 representing “No image at all, you only ‘know’ that you are thinking of the object”. Lower scores on VVIQ indicate higher mental imagery ability on visual imagery vividness. This measure shows high internal reliability with a split-half reliability of .85 in the original paper. In the current study, the Cronbach’s alpha was .95.

### Analysis

Because of technical difficulty during data collection, some participants were excluded from analysis for a specific measure. The final sample sizes were as follows: *n* = 54 for EMGc and EMGz; *n* = 54 for HR; *n* = 59 for EDA and SKT; *n* = 52 for BAIS-V.

To examine the effect of aversiveness across modalities, we conducted a two-factor within-subjects ANOVA. *A priori* t-tests were conducted to assess the aversiveness effect within each of the three modalities (H1 and H2). Bonferroni correction was applied for any post-hoc tests. For each physiological measure, any trial that had a value larger than 3 standard deviations was considered as an outlier and removed from the subsequent analyses. The percentages of the trials that were removed were 2.02% for EMGc; 1.98% for EMGz; 0.65% for EDA; 0.83% for HR; 1.14% for SKT.

To examine the similarity of the physiological response patterns in aversiveness differentiation across modalities (H3), we trained a support vector machine (SVM) classifier with the radial based function kernel on the facial and autonomic signals of one modality and tested on another modality. Specifically, each of the five physiological features (EMGc, EMGz, EDA, HR and SKT) was averaged across trials separately by aversiveness and modality for every participant. The missing values were replaced by the mean value across participants per aversiveness and modality, resulting in a sample consisting of all participants (*N* = 59). Those collapsed features were used to classify the aversiveness (aversive vs. nonaversive). The accuracy score was reported to evaluate the cross-modality classification performance. The aversiveness labels of the training set were randomly shuffled and the same cross-modality model was run 1,000 times to obtain a permutation distribution of the classification accuracy. The 95th percentile of that accuracy distribution served as the critical value for the accuracy significantly higher than chance level (*p* < .05, one-tailed). The by-modality classification, training and testing the classifier on the same modality, was also conducted serving as a comparison baseline. A 10-fold cross-validation was applied, such that the data from the same participant was not part of both training and testing sets. The permutation test as mentioned above was applied to obtain the critical value of accuracy for different modalities.

To test the effect of vividness of auditory imagery on behavioral ratings and physiological responses (H4), multiple linear regression analyses were conducted. Specifically, for each of the behavioral and physiological responses, we computed difference scores by subtracting the average value of the responses to nonaversive trials from the aversive trials for the imagery modality. Next, we used the BAIS-V scores to predict difference scores, controlling for VVIQ, age, and sex.

## Results

### Manipulation of aversiveness

We used 20 aversive and 20 nonaversive stimuli across three different modalities to manipulate participants’ behavioral and physiological responses. Our stimuli resulted in an expected separation in aversiveness perception based on the average valence ratings (Figure 2A). All aversive stimuli had average valence ratings of at most 3 (on a 0-9 scale, where 0 represents very negative and 9 represents very positive), with the exception of the audiovisual and imagery trials of one stimulus (“Someone squishing wet clothes”, see Figure 2A). Most nonaversive stimuli had average valence ratings of at least 4, with the exception of the auditory trials for three stimuli (“Sizzling in a frying pan”, “A water sprinkler”, and “Pouring uncooked rice”). Although arousal ratings were fairly similar across aversiveness, they were significantly higher for aversive compared to nonaversive stimuli (Mean = 5.89 for aversive stimuli, Mean = 5.33 for nonaversive stimuli), *t*(38) = 3.14, *p* < .01, Cohen’s *d* = 0.99.

**Figure 2.**
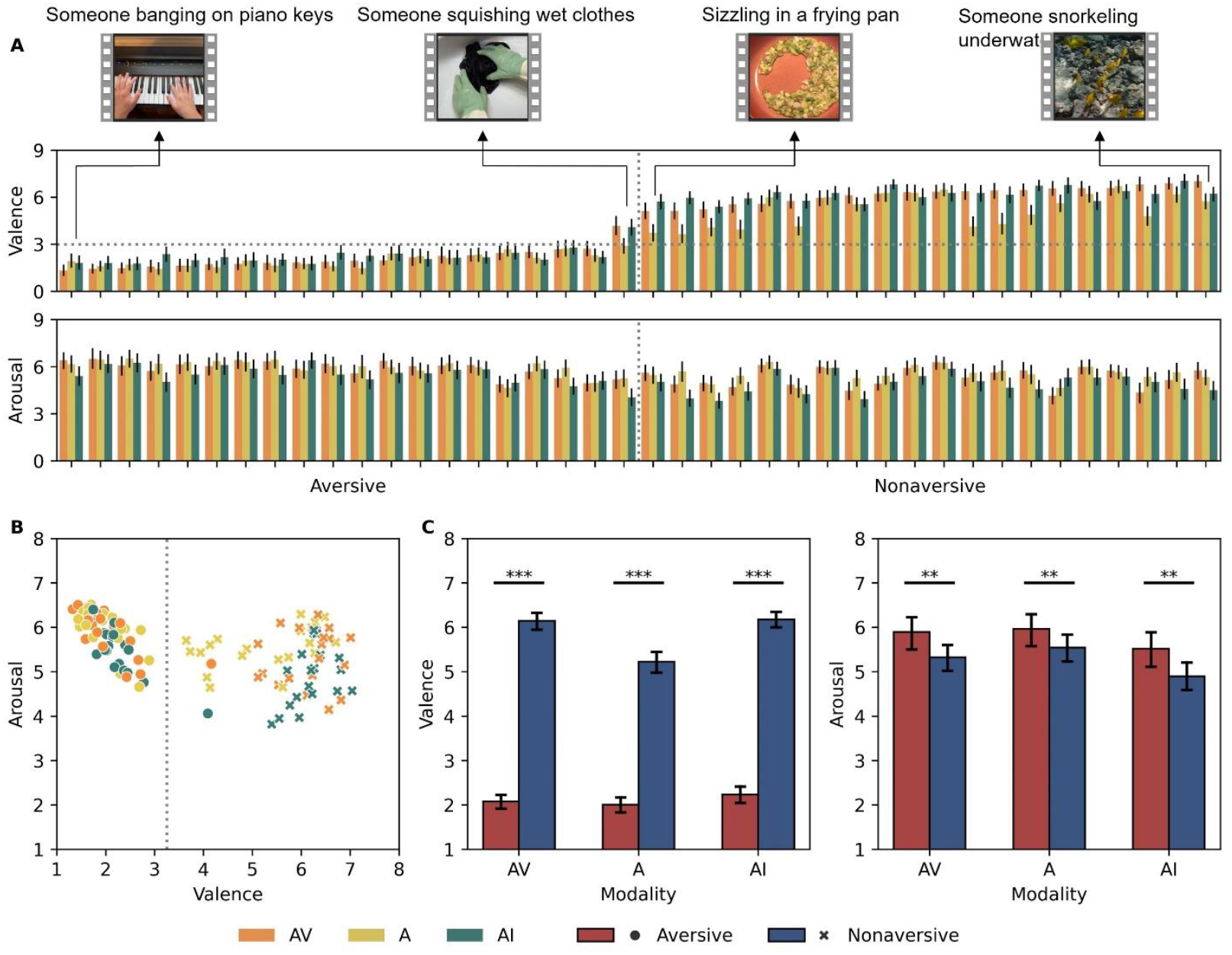
**A**. Average valence and arousal ratings by modality for the 40 stimuli used in this study, ordered by the valence ratings on AV stimuli from the lowest to the highest. **B**. The bivariate scatter plot between valence and arousal ratings by modality for the 40 stimuli used in this study. The dotted line denotes a separation of aversiveness. **C**. Aversive and nonaversive trials were well separated on valence for each modality. Aversive and nonaversive trials also differed on arousal. *Notes*. ^**^*p* <. 01, \*\*\**p* < .001. Error bars denote 95% confidence interval.

### Valence and arousal ratings

For valence ratings, we found a significant main effect of modality, *F*(2,116) = 65.96, *MSE* = 0.18, 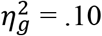, a significant main effect of aversiveness, *F*(1,58) = 684.86, *MSE* = 1.81, 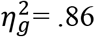, and a significant interaction effect, *F*(2,116) = 30.45, *MSE* = 0.20, *p* <.001, 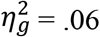 (Figure 2). Planned t-tests showed significant difference between the valence ratings for aversive and nonaversive stimuli for AV, mean difference = −4.07, 95% CI = [−4.35, −3.79], *t*(58) = −28.83, *p* < .001, Cohen’s *d* = 3.75, A, mean difference = −3.22, 95% CI = [−3.56, −2.89], *t*(58) = −18.94, *p* < .001, Cohen’s *d* = 2.47, and AI, mean difference = −3.94, 95% CI = [−4.26, −3.62], *t*(58) = −24.34, *p* < .001, Cohen’s *d* = 3.17. After Bonferroni correction, the post-hoc comparisons showed that the valence ratings were significantly lower for A compared to AV, mean difference = −0.49, 95% CI = [−0.64, −0.35], *t*(58) = −8.32, *p* < .001, Cohen’s *d* = 0.63 and AI, mean difference = −0.59, 95% CI = [−0.73, −0.45], *t*(58) = −10.10, *p* < .001, Cohen’s *d* = 0.76. There was no significant difference between AV and A.

For arousal ratings, we found significant main effects of modality, *F*(2,116) = 8.62, MSE = 1.10, 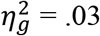 and aversiveness, *F*(1,58) = 10.06, *MSE* = 2.55, 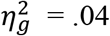 (Figure 2).

The interaction effect was not significant. After Bonferroni correction, the post-hoc t-tests showed significant differences between the arousal ratings for aversive and nonaversive stimuli for AV, mean difference = 0.57, 95% CI = [0.16, 0.98], *t*(58) = 2.74, *p* = .024, Cohen’s *d* = 0.36, A, mean difference = 0.42, 95% CI = [0.15, 0.70], *t*(58) = 3.04, *p* = .012, Cohen’s *d* = 0.40, and AI, mean difference = 0.62, 95% Ci = [0.23, 1.02], *t*(58) = 3.13, *p* < .01, Cohen’s *d* = 0.41. The arousal ratings were significantly higher for AV, mean difference = 0.40, 95% CI = [0.18, 0.62], *t*(58) = 2.59, *p* = .036, Cohen’s *d* = 0.33, and A compared to AI, mean difference = 0.55, 95% CI = [0.30, 0.79], *t*(58) = 3.37, *p* < .01, Cohen’s *d* = 0.40.

### Physiological responses

For EMGc, our planned comparisons showed significant greater potentiation for aversive AV, mean difference = 0.20, 95% CI = [0.09, 0.30], *t*(53) = 3.61, *p* < .001, Cohen’s *d* = 0.49, A, mean difference = 0.15, 95% CI = [0.05, 0.25], *t*(53) = 3.02, *p* < .01, Cohen’s *d* = 0.41, and AI stimuli, mean difference = 0.07, 95% CI = [0.02, 0.13], *t*(53) = 2.56, *p* = .013, Cohen’s *d* = 0.35, compared to nonaversive stimuli (Figure 3). The two-way repeated measures ANOVA showed a significant main effect of modality, *F*(2,106) = 8.13, *MSE* = 0.08, *p* < .001, 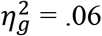, a significant main effect of aversiveness, *F*(1,53) = 12.36, *MSE* = 0.13, *p* < .001, 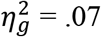, and a significant interaction effect, *F*(2,106) = 3.33, *MSE* = 0.06, *p* = .040,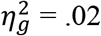. After Bonferroni correction, the post-hoc comparisons showed that the EMGc potentiation was significantly greater for AV compared to AI, mean difference = 0.15, 95% CI = [0.08, 0.21], *t*(53) = 4.39, *p* < .001, Cohen’s *d* = 0.45. There were no significant differences between AV and A or A and AI.

**Figure 3.**
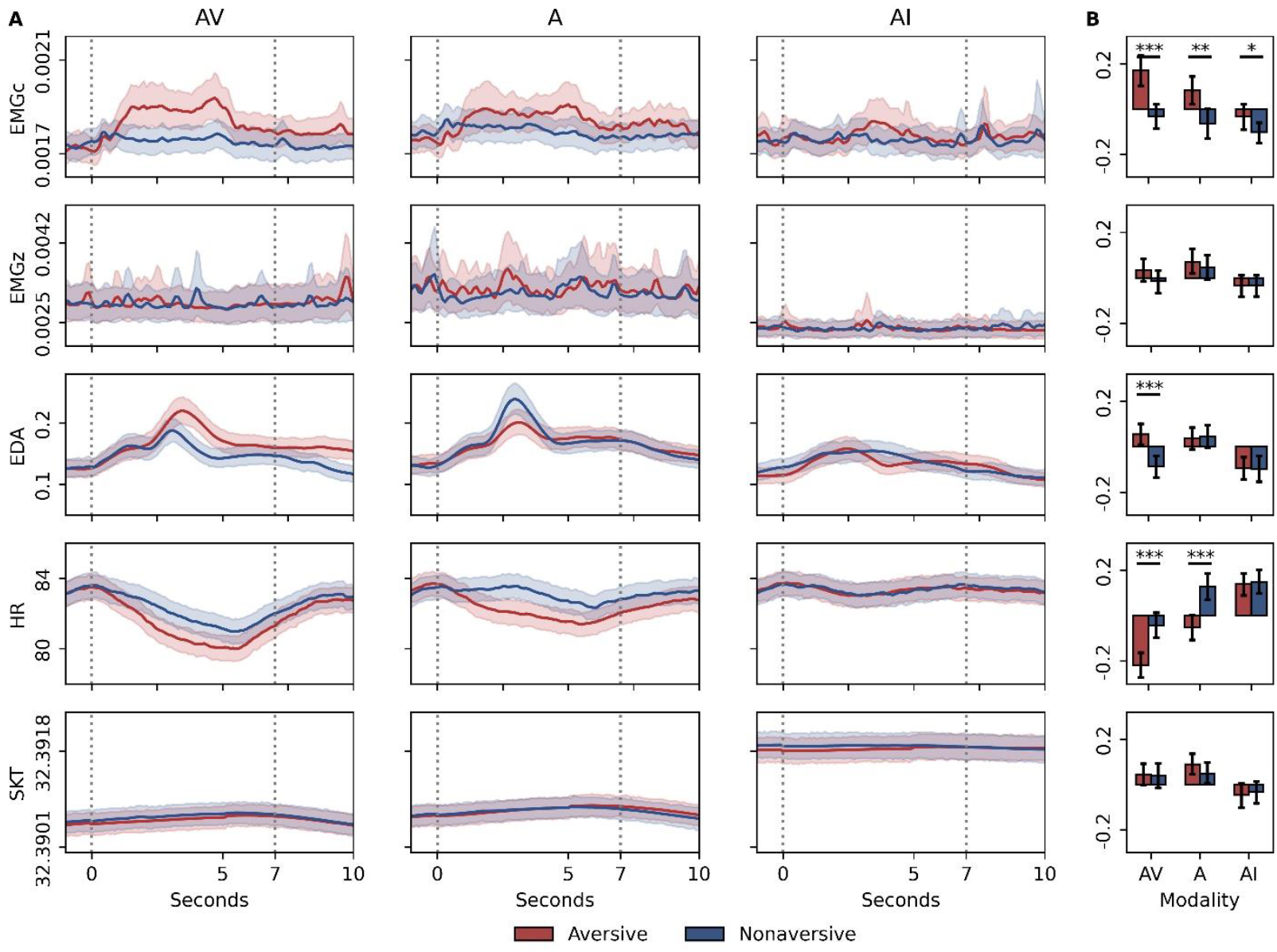
**A**. Visualization of the raw signal changes across time. The signals were averaged across trials and participants. Dotted lines mark the 7 s time window following stimulus presentation used for data analysis. Shaded regions denote 95% confidence interval. **B**. The aversiveness effect by modality across physiological measures (z-scores). Error bars denote 95% confidence interval. *Notes*. ^*^*p* < .05, ^**^*p* <. 01, ^***^*p* < .001.

For EMGz, our planned comparisons did not show significant aversiveness effects for any of the three modalities (Figure 3). The two-way repeated measures ANOVA showed a significant modality effect, *F*(2,106) = 3.80, *MSE* = 0.07, *p* = .025,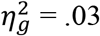. The main effect of aversiveness and the interaction effect were not significant. After Bonferroni correction, the post-hoc comparisons for the modality effect only showed larger potentiation for A compared to AI, mean difference = 0.09, 95% CI = [0.03, 0.15], *t*(53) = 2.67, *p* = .030, Cohen’s *d* = 0.29. There were no significant differences between AV and A or AV and AI.

For EDA, the planned comparison showed significant greater skin conductance level for aversive AV stimuli compared to nonaversive AV stimuli, mean difference = 0.14, 95% CI = [0.08, 0.20], *t*(53) = 4.11, *p* < .001, Cohen’s *d* = 0.54 (Figure 3). The aversiveness effect was not significant for A or AI. The two-way repeated measures ANOVA showed a significant main effect of modality, *F*(2,116) = 12.67, *MSE* = 0.04, *p* < .001, 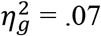, a significant main effect of aversiveness, *F*(1,58) = 4.03, *MSE* = 0.04, *p* = .049, 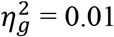, and a significant interaction effect, *F*(2,116) = 5.12, *MSE* = 0.05, *p* < .01,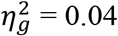. After Bonferroni correction, the post-hoc comparisons showed that EDA was significantly greater for A, mean difference = 0.14, 95% CI = [0.08, 0.20], *t*(58) = 4.71, *p* < .001, Cohen’s *d* = 0.43, and AV, mean difference = 0.08, 95% CI = [0.02, 0.14], *t*(58) = 2.78, *p* = .022, Cohen’s *d* = 0.26, compared to AI. There was no significant difference between A and AV.

For HR, the planned comparisons showed significant heart rate deceleration for aversive AV, mean difference = −0.16, 95% CI = [−0.24, −0.08], *t*(53) = −4.15, *p* < .001, Cohen’s *d* = 0.56, and A, mean difference = −0.19, 95% CI = [−0.28, −0.10], *t*(53) = −4.24, *p* < .001, Cohen’s *d* = 0.58, but not for AI compared to nonaversive stimuli (Figure 3). The two-way repeated measures ANOVA showed a significant main effect of modality, *F*(2,106) = 32.80, *MSE* = 0.07, *p* < .001, 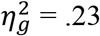, a significant main effect of aversiveness, *F*(1,53) = 36.84, *MSE* = 0.04, *p* < .001, 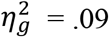, and a significant interaction effect, *F*(2,106) = 5.29, *MSE* = 0.05, *p* < .01,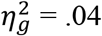. After Bonferroni correction, the post-hoc comparisons showed significantly larger heart rate deceleration for AV compared to A, mean difference = −0.17, 95% CI = [−0.23, −0.10], *t*(53) = −5.15, *p* < .001, Cohen’s *d* = 0.50, and AI, mean difference = −0.29, 95% CI = [−0.35, −0.22], *t*(53) = −8.38, *p* < .001, Cohen’s *d* = 0.88. The heart rate deceleration for A was significantly larger than that for AI, mean difference = −0.12, 95% CI = [−0.18, −0.06], *t*(53) = −3.92, *p* < .001, Cohen’s *d* = 0.36.

For SKT, our planned comparisons did not show significant aversiveness effects for any of the three modalities (Figure 3). The two-way repeated measures ANOVA only showed a significant modality effect, *F*(2,116) = 11.20, *MSE* = 0.04, *p* < .001,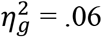. The main effect of aversiveness and the interaction effect were not significant. After Bonferroni correction, the post-hoc comparisons showed that the finger SKT change for AI was significantly lower than that for AV, mean difference = −0.08, 95% CI = [−0.14, −0.02], *t*(53) = −3.13, *p* < .01, Cohen’s *d* = 0.25, and A, mean difference = −0.11, 95% CI = [−0.16, −0.06], *t*(53) = −4.39, *p* < .001, Cohen’s *d* = 0.39. There was no significant difference between AV and A.

### The classification of aversiveness based on physiological responses

The by-modality results showed that the accuracy of aversiveness classification based on the response pattern in EMGc, EMGz, EDA, HR, and SKT was significantly higher than chance level for AV stimuli, but not for A or AI stimuli (Table 1).

**Table 1.** Accuracy of the aversiveness classification by modality and cross modality.

| Model | Classification accuracy | Threshold |
| --- | --- | --- |
| By-modality |  |  |
| AV | .702* | .585 |
| A | .577 | .585 |
| AI | .515 | .585 |
| Cross-modality |  |  |
| Train on AV, test on A | .669* | .568 |
| Train on AV, test on AI | .517 | .534 |
| Train on A, test on AV | .619* | .568 |
| Train on A, test on AI | .492 | .559 |
| Train on AI, test on AV | .636* | .568 |
| Train on AI, test on A | .585 | .585 |

Note. \* $p < .05$ .

The cross-modality results revealed a similar physiological response pattern between AV and A in differentiating aversive and nonaversive affective states. The physiological response pattern for AI was useful in classifying aversiveness for AV but not for A. However, the accuracy of using the physiological pattern found in AV or A was not significantly better than chance level when predicting aversiveness for AI.

### The effect of auditory imagery vividness

Multiple linear regression analyses were conducted to test the effect of vividness of auditory imagery on behavioral ratings and physiological responses while participants were prompted to imagine sounds. The model was significant for difference in valence ratings, *F*(4,46) = 3.05, *p* = .026, adjusted R squared = .14. Controlling for VVIQ, age, and sex, the BAIS-V score was negatively correlated with the difference in valence ratings between aversive and nonaversive stimuli, *β* = −.03, *SE* = .012, *p* = .013. Participants with more vivid auditory imagery had larger separation in valence ratings between imagined aversive and nonaversive sounds. No significant results were found for differences in arousal ratings or physiological responses.

## Discussion

The major goal of the current study was to examine somatovisceral responses when imagining aversive and nonaversive naturalistic sounds. We compared the similarity of the facial and autonomic responses under aversive and nonaversive affective states across audiovisual, auditory, and auditory imagery modalities. In addition, we also assessed the effect of individual differences in auditory imagery vividness on the subjective ratings and physiological responses.

Our results replicated previous findings that exogenously induced aversive affective state correspond to coherent changes in subjective experience, facial expression, and electrodermal and cardiac activities for both audiovisual (H1a) and auditory modalities (H1b). Compared to nonaversive stimuli, aversive audiovisual and auditory stimuli both induced EMG potentiation over the corrugator and sustained heart rate deceleration. There was a significant phasic EDA increase for aversive audiovisual stimuli, but not for aversive auditory stimuli. These results were in line with previous evidence using affective films (Golland et al., 2018; Sato et al., 2020; Sato & Kochiyama, 2022) and affective sounds (Bradley & Lang, 2000; Larsen et al., 2003).

Contrary to our hypothesis, the results did not show significant differentiation between aversive and nonaversive stimuli for EMGz, supporting previous evidence that EMGc has a closer correspondence to aversiveness as opposed to EMGz (Larsen et al., 2003). This lack of differentiation for EMGz cannot be explained by the quadratic relationship hypothesis that previously showed that EMGz potentiates for both extreme aversive and nonaversive stimuli (Bradley & Lang, 2000; Lang et al., 1993; Larsen et al., 2003). In the current study, the EMGz activity was weak, rather than strong, for both aversive and nonaversive stimuli. One possible explanation is that, as addressed by Larsen (2003), there is a “threshold effect” for the EMGz responses. Specifically, EMG activity in this muscle site can only be observed when people process extreme valenced stimuli exceeding a valence threshold. The stimuli used in the current study do not contain extreme affective content and thus may be insensitive to EMG zygomaticus responses. In addition, we did not find any significant aversiveness differentiation for finger SKT. As suggested by previous evidence, SKT indexes the arousal change between highly arousing aversive and nonaversive stimuli and less arousing neutral stimuli (Ioannou et al., 2014; Sato et al., 2020; Sato & Kochiyama, 2022). Future studies may include neutral stimuli and extend the aversiveness range to obtain a better understanding of these two physiological measures showing null results.

The major goal of the current study was to examine the physiological responses to the auditory imagery of aversive and nonaversive sounds. Our results revealed significant differentiation in subjective hedonic valence ratings and EMGc potentiation between aversive and nonaversive states during auditory imagery (H2). Auditory imagery vividness predicted larger separation in subjective ratings on hedonic valence between aversive and nonaversive auditory images, but not the physiological responses (H4). Autonomic activity, including electrodermal, cardiac, and skin temperature responses, were generally weaker as compared to previous evidence on affective visual imagery reporting coherent changes across facial and autonomic measures in which aversive imagery indexes potentiated EMG corrugator and skin conductance activity and accelerated heart rate (Bradley & Lang, 2007). There are several potential explanations. First, early studies have shown that imagery scripts play a critical role in the success of the manipulation of physiological activation. Our auditory imagery prompts (e.g., imagine the sound of “someone banging on piano keys”) were stimulus-proposition-laden and may have been too concise to induce auditory images carrying strong affective content. Most of the studies investigating affective imagery use long detailed scripts containing rich information (Acosta & Vila, 1990; Lang et al., 1980; G. A. Miller et al., 1987). Lang et al. (1980) found that a stimulus-proposition-laden script (e.g., “you are flying a kite on the beach on a bright summer day”) failed to elicit appreciable physiological responses during affective imagery, whereas a response-proposition-laden scripts (e.g., “you breathe deeply as you run along the beach flying a kite”) elicited a significant reaction pattern in EMG, HR and skin conductance level.

Second, the imagining period in our study was relatively short (5 s) compared to previous studies asking the participants to imagine a scene in a longer time frame ranging from 30 s to several minutes (Acosta & Vila, 1990; Lang et al., 1980; Schwartz et al., 1976, 1980). Five seconds may not be long enough for the participants to generate and engage in the imagery. Some of our imagery prompts require extra effort to retrieve the sounds from memory first (e.g., “Upbeat marching band music” and “A teapot whistling”). Thus, a shorter time window may not be sufficient to elicit a strong aversive affective state.

A final explanation is that the weak results across autonomic measures might reflect a true modality effect of auditory imagery. In our data, we found that the aversive auditory imagery did elicit a more aversive subjective experience (reflected by valence ratings) but failed to elicit significant physiological responses (except for EMGc). Engen et al. (2018) found that auditory imagery showed poor efficacy in generating emotions as compared to visual imagery and bodily interoception. Future work is needed to tease out these possible explanations through engineering the imagery script and manipulating the imagery context.

To examine the similarity in physiological response patterns across modalities, we tested if a classifier trained on one modality can predict the aversive and nonaversive states of another modality. This would support modality general physiological representation of affect (H3). Our results revealed that a simple SVM classifier trained on the somatovisceral responses to aversive and nonaversive audiovisual stimuli without feature engineering and hyperparameter tuning is useful to predict the aversiveness of auditory modality, and vice versa, suggesting a similar physiological reaction pattern underlying aversiveness perception across sensory modalities. Similar results were reported by Kim and Wedell (2016), who found that the physiological mechanism underlying valence processing was represented in a similar way across visual (pictures) and auditory (music) modalities. Interestingly, the models trained on audiovisual and auditory modalities failed to predict the aversiveness of auditory imagery, but the physiological pattern during affective auditory imagery was useful in predicting the aversiveness of audiovisual stimuli, likely driven by EMGc potentiation. However, this result should be interpreted with caution, because a model trained on noisy data (auditory imagery) may successfully learn the features that can be generalized to less noisy data (audiovisual) but not the other way around.

## Conclusions

We examined somatovisceral responses to aversive and nonaversive stimuli across three modalities: audiovisual, auditory, and auditory imagery. The results revealed that EMGc potentiation for aversive stimuli was the most coherent response across modalities. Physiological responses to the auditory imagery of aversive and nonaversive sounds prompted by short text-based scripts were weaker compared to responses elicited by audiovisual and auditory stimuli.

The cross-modality classification results showed an underlying similarity in the physiological response patterns between audiovisual, auditory, and auditory imagery conditions. The current study has important implications in understanding the physiological mechanisms underlying affective auditory imagery.

## Acknowledgement

Thanks COR for helpful discussion. This research was supported by a grant from the Misophonia Research Fund.

